# Distribution of Huntington’s disease Haplogroups in Indian population

**DOI:** 10.1101/2020.12.31.424975

**Authors:** Sowmya Devatha Venkatesh, Nikhil Ratna, Swathi Lakshmi.P, Geetanjali Murari, Nitish Kamble, Ravi Yadav, Biju Viswanath, Mathew Varghese, Pramod Kumar Pal, Sanjeev Jain, Meera Purushottam

## Abstract

Huntington’s disease (HD), a rare neurodegenerative disorder, is inherited in an autosomal dominant manner, and caused by a pathological trinucleotide expansion at exon1 of the *HTT* locus. Previous studies have described the haplogroups at the *HTT* locus that can explain the differences in prevalence of HD. We have selected three informative SNPs (rs762855, rs3856973 and rs4690073) to study these haplogroups in an Indian sample. Our results show that the genotype frequencies are significantly different between cases and controls for these SNPs. More than 90% of both cases and controls belong to Haplogroup A which is the predominant European haplogroup.

## Introduction

Huntington’s disease (HD) is an autosomal dominant, neurodegenerative disorder, caused by an expansion of an unstable CAG triplet repeat sequence in the Huntingtin (*HTT*) gene. The size of the repeats varies from 17-20, in most individuals. Expansion of this CAG repeat stretch upto 26, is thought not to contribute to disease, while CAG repeats above 36 are considered as disease causing. Correspondingly, CAG repeats from 27 to 35 are considered to be intermediate alleles. The prevalence of HD varies across the world, by both geography (prevalence 5-15/100,000 in Europe; and 1-2/100,000 in sub-Saharan Africa and China); and ethnicity. The number of CAG repeats in the normal allele, and the frequency of intermediate alleles (IAs) perhaps has a bearing on prevalence. One study on the European and American populations showed the intermediate alleles ranged from 0.45% - 8.7% in the general population, 0.05% - 5.1% in the individuals with family history of HD; and was highest in the general Brazilian population (8.7%)^(1)^. Another study in the Northern part of Sweden (SHAPE; The Swedish Huntingtin Alleles and Phenotype) observed that 6.3% of 7379 individuals from the general population had IAs, and there were even some individuals with very short (<5) with repeat numbers ^(2)^. CAG repeat sizing of polyglutamine disorders in the general population showed the HD locus to have a high proportion of IAs; 6% in European (N=13,670)^(3)^ and 5.3% in Italian populations (N= 729)^(4)^.

Haplotype studies on the *HTT* locus have identified three major haplogroups (A, B and C) which differ across populations^(1, 6)^. Haplogroup A (subgroups A1 & A2), B and C are the major haplogroups of European, African and East Asian HD cases respectively^(5–7)^. There is a similar pattern of haplogroup representation in the IAs, as in HD cases^(8)^, suggesting that IAs with the high risk haplogroups are more likely to transmit HD, compared to others. If alleles with certain haplotypes are more susceptible to expansions than others, tracking these may help us understand the factors that contribute to differences in prevalence, and outcomes. Our study thus attempts to understand the haplogroup structure in persons with Huntington’s disease from India.

## Materials & Methods

We have taken HD (N 196; F 74) and healthy controls (N 200; F 88) for the study. All the HD patients were identified in the clinical services, and tested at the Genetic Counseling and Testing (GCAT) clinic of National Institute of Mental Health and Neurosciences (NIMHANS). Healthy controls are volunteers with no neurological or psychiatric illness. Informed consent has been taken for both HD cases and healthy controls. The study was carried out after institutional ethical clearance. Genomic DNA was extracted from peripheral blood and PCR was done for CAG allele determination for both cases and controls with specific primers within the *HTT* gene^(9)^. The three SNPs located at the *HTT* locus on chromosome 4 (rs ID/ gnomAD variant ID: rs762855/4-3074794-A-G, rs3856973/4-3080173-G-A and rs4690073/4-3160150-G-A) were also genotyped. These three SNPs selected from the 22 tag SNPs described in a previous study (3), were used to distinguish between these haplogroups in those detected to have HD and healthy controls. High resolution melt curve (HRM) analysis was done for rs762855 and PCR-RFLP method for rs3856973 and rs4690073. Validation was done by Sanger sequencing. The primer sequences used for PCR amplification are: rs762855 FP - GCAGTAGCCTCCCTTTTCTTG, RP - TCAAATTCCTGGGTTCAGGT; rs3856973 FP - CAGCAGTGAGCAGACAAAGC, RP - TTCTGGGTTTTGCTGGAAAG and rs4690073 FP - GGGATCAGTTCCCCTGTTGT, RP - CCATGCAGCTTAAAGAGACCT. Haplotype construction was done using homozygous genotypes for the three SNPs (only homozygous genotypes were selected for haplotype construction as heterozygotes could not be assigned to haplotypes unambiguously). The haplogroups are ascribed based on allele combination of SNPs as Haplogroup A - AGG, Haplogroup B - GGG and Haplogroup C-GAA corresponding to the SNP sequence rs762855, rs3856973 followed by rs4690073. The available phased 1000 genome data of the three SNPs was taken for the south Asian population and haplogroups were constructed (Fig 1).

**Fig I:**
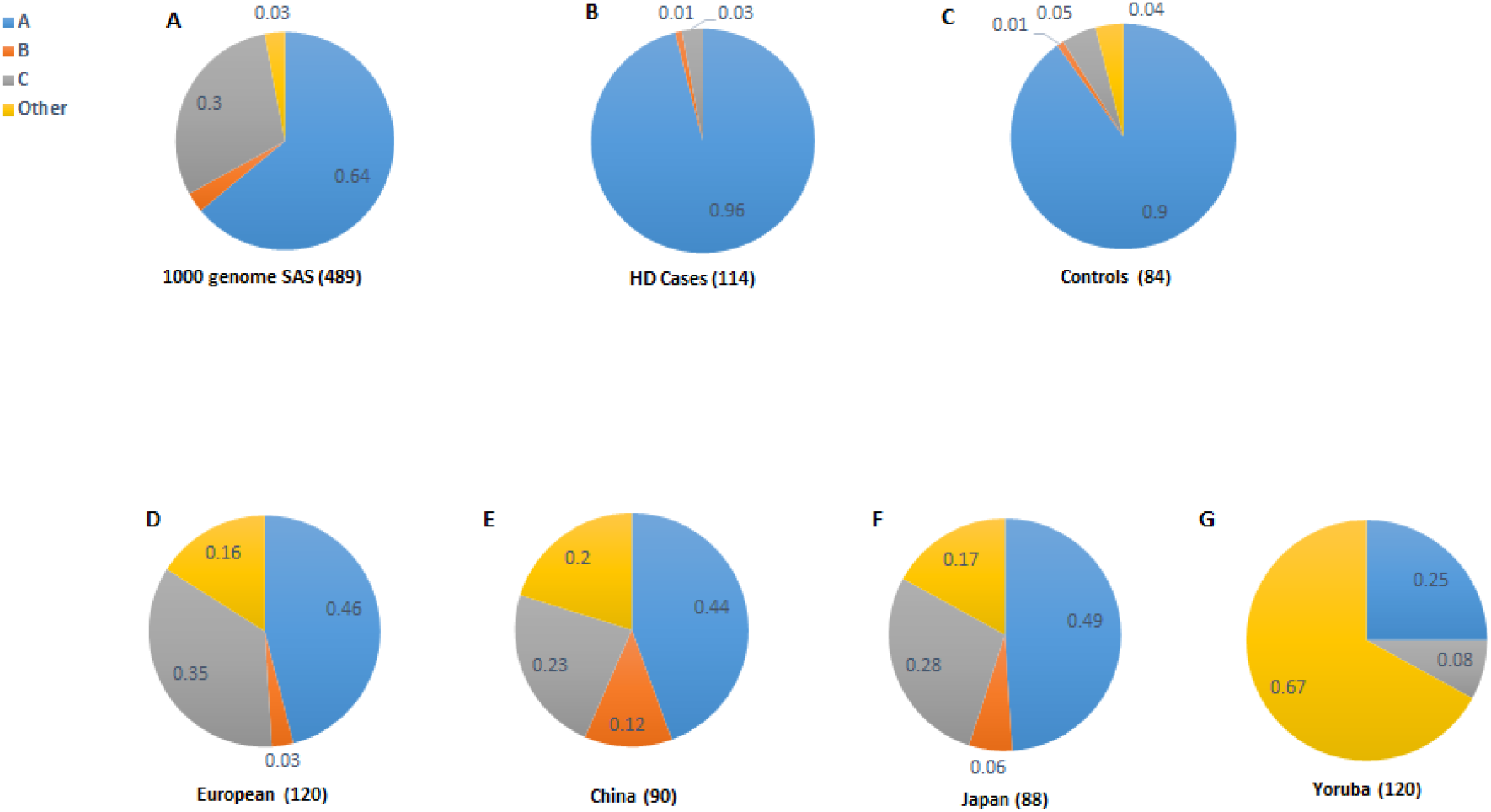
Haplogroup distribution comparison of our study with other populations. The haplogroup distribution shown in percentage, A - 1000 genome data, B and C (homozygous genotypes were selected for haplogroup construction). - HD cases and controls from our study, D, E, F and G - European, China, Japan and Yoruba general population, adapted from (5)

Comparison of genotype frequency between cases and controls was done by Chi-square test.

## Results & Discussion

The CAG repeat allele distribution at the *HTT* locus for HD patients (normal allele - 17.72±2.55 (12-32), expanded allele - 45.64±7.95 (39-113)) and controls (normal allele1 - 16.6±1.35 (10-22), normal allele 2 - 18.8± 2.7 (14-29)) was determined. An intermediate allele was detected in [N= 3, 1.4%] controls; and in [N=3, 1.5%] HD patients with an expanded HD allele, where the other allele was in the intermediate range. The distribution of three SNPs followed Hardy-Weinberg equilibrium, for both cases and controls.

The allele and genotype frequencies of the three selected SNPs were significantly different between HD cases and controls (Table I). The haplotype could be unambiguously ascertained in 203 persons (119 HD; 84 controls); and in these the predominant haplogroup was Haplogroup A (114 cases; 76 controls). On the other hand Haplogroup B (1 each in cases and controls) and C (4 each in cases and controls) were relatively rare. The frequency distribution of haplogroups is shown in Figure I. All the three SNPs were in strong linkage disequilibrium.

**Table I:**
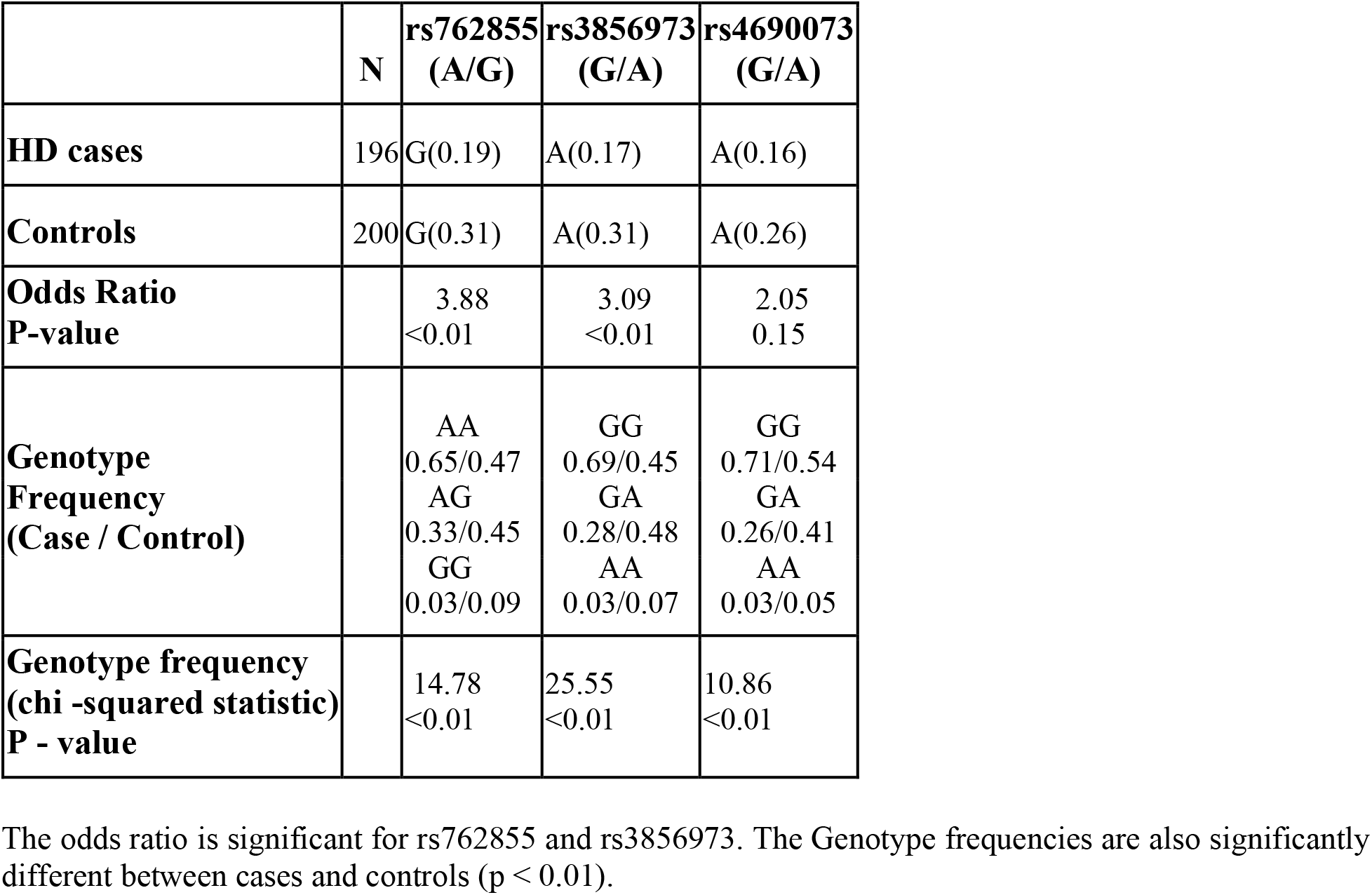
Allele and genotype frequencies of the 3 studied SNPs.

We detect that the allele frequencies of three SNPs studied at *HTT* loci differ in subjects with HD, when compared to the background population, as expected. The predominant haplogroup, as defined by these SNPs, however, was the Haplogroup A, in HD patients, as has been reported earlier. However, a few individuals do conform to haplogroup C. This hints that there may be multiple founder events, as well as admixture, that contributes to this diversity in India.

The identified haplogroup broadly overlaps with that of the European population, however it should be borne in mind that variants of Haplogroup A have also been seen in the East Asian and African populations. Since only a subset of samples were homozygous, this description may be valid for only a subset of the individuals. A study with more SNPs/markers around the *HTT* gene locus to aid better classification into haplogroups would help us obtain a better idea about the origin and spread of the disease, and also plan future interventions, such as allele specific therapeutics. Such knowledge could also have a bearing on the underlying mechanisms of other neuro-psychiatric diseases, especially those associated with unstable repeat sequences.

## Acknowledgment

The authors would like to thank the Genetic Counseling and Testing (GCAT) clinic and all the patients, their family members and volunteers for their participation and cooperation.

## Financial support & Sponsorship

Authors would like to thank DST-SERB (India) for the financial support provided for this work.

## Conflict of Interest

The authors have no conflict of interest to report.

## Notes

### Competing Interest Statement

The authors have declared no competing interest.

## References

1. Apolinário TA, Paiva CLA, Agostinho LA. REVIEW-ARTICLE Intermediate alleles of Huntington’s disease HTT gene in different populations worldwide: a systematic review. Genet Mol Res. 2017 Apr 5;16(2).

2. Sundblom J, Niemelä V, Ghazarian M, Strand A-S, Bergdahl IA, Jansson J-H, et al. High frequency of intermediary alleles in the HTT gene in Northern Sweden - The Swedish Huntingtin Alleles and Phenotype (SHAPE) study. Scientific Reports. 2020 Jun 17;10(1):9853.

3. Gardiner SL, Boogaard MW, Trompet S, de Mutsert R, Rosendaal FR, Gussekloo J, et al. Prevalence of Carriers of Intermediate and Pathological Polyglutamine Disease–Associated Alleles Among Large Population-Based Cohorts. JAMA Neurol. 2019 Jun 1;76(6):650.

4. Mongelli A, Magri S, Salvatore E, Rizzo E, De Rosa A, Fico T, et al. Frequency and distribution of polyQ disease intermediate-length repeat alleles in healthy Italian population. Neurol Sci. 2020 Jun 1;41(6):1475–82.

5. Warby SC, Montpetit A, Hayden AR, Carroll JB, Butland SL, Visscher H, et al. CAG Expansion in the Huntington Disease Gene Is Associated with a Specific and Targetable Predisposing Haplogroup. The American Journal of Human Genetics. 2009 Mar;84(3):351–66.

6. Baine FK, Kay C, Ketelaar ME, Collins JA, Semaka A, Doty CN, et al. Huntington disease in the South African population occurs on diverse and ethnically distinct genetic haplotypes. Eur J Hum Genet. 2013 Oct;21(10):1120–7.

7. Warby SC, Visscher H, Collins JA, Doty CN, Carter C, Butland SL, et al. HTT haplotypes contribute to differences in Huntington disease prevalence between Europe and East Asia. Eur J Hum Genet. 2011 May;19(5):561–6.

8. Kay C, Collins JA, Wright GEB, Baine F, Miedzybrodzka Z, Aminkeng F, et al. The molecular epidemiology of Huntington disease is related to intermediate allele frequency and haplotype in the general population. Am J Med Genet B Neuropsychiatr Genet. 2018;177(3):346–57.

9. Warner JP, Barron LH, Brock DJH. A new polymerase chain reaction (PCR) assay for the trinucleotide repeat that is unstable and expanded on Huntington’s disease chromosomes. Molecular and Cellular Probes. 1993 Jun;7(3):235–9.

